# Chromatin structure and *var2csa* – a tango in regulation of *var* gene expression in the human malaria parasite, *Plasmodium falciparum*?

**DOI:** 10.1101/2024.02.13.580059

**Authors:** Todd Lenz, Xu Zhang, Abhijit Chakraborty, Abbas Roayaei Ardakany, Jacques Prudhomme, Ferhat Ay, Kirk Deitsch, Karine G. Le Roch

## Abstract

Over the last few decades, novel methods have been developed to study how chromosome positioning within the nucleus may play a role in gene regulation. Adaptation of these methods in the human malaria parasite, *Plasmodium falciparum*, has recently led to the discovery that the three-dimensional structure of chromatin within the nucleus may be critical in controlling expression of virulence genes (*var* genes). Recent work has implicated an unusual, highly conserved *var* gene called *var2csa* in contributing to coordinated transcriptional switching, however how this gene functions in this capacity is unknown. To further understand how *var2csa* influences *var* gene switching, targeted DNA double-strand breaks (DSBs) within the sub-telomeric region of chromosome 12 were used to delete the gene and the surrounding chromosomal region. To characterize the changes in chromatin architecture stemming from this deletion and how these changes could affect *var* gene expression, we used a combination of RNA-seq, Chip-seq and Hi-C to pinpoint epigenetic and chromatin structural modifications in regions of differential gene expression. We observed a net gain of interactions in sub-telomeric regions and internal *var* gene regions following *var2csa* knockout, indicating an increase of tightly controlled heterochromatin structures. Our results suggest that disruption of *var2csa* results not only in changes in *var* gene transcriptional regulation but also a significant tightening of heterochromatin clusters thereby disrupting coordinated activation of *var* genes throughout the genome. Altogether our result confirms a strong link between the *var2csa* locus, chromatin structure and *var* gene expression.

**AUTHOR SUMMARY:** Malaria remains one of the deadliest parasite-borne diseases, causing not only over a half million deaths annually, but also infecting hundreds of millions more. *Plasmodium falciparum,* the protozoan parasite that is responsible for the most virulent form of human malaria, is transmitted to humans by infected female mosquitoes during a blood meal. Due to a growing resistance to all existing antimalarials, there is a need to identify novel targets to design new antimalarial strategies. Our research builds on the growing body of evidence that supports the role of genome organization or chromatin structure within the nucleus in controlling the parasite development as well as virulence factors designed to circumvent the host immune response. This study identifies genes and structural elements within the *Plasmodium falciparum* genome that are controlled, at least partially, by the expression of a single unique and highly conserved virulence gene.

## INTRODUCTION

Although the recent coronavirus pandemic has dominated the news for the last few years, it is important to note that human malaria parasites still infect 249 million people worldwide annually. *Plasmodium*, the causative agent of malaria, remains one of the deadliest parasite-borne diseases, with an estimated 608,000 deaths in 2022 [1] affecting mostly children and immunocompromised individuals (e.g., pregnant women and the elderly) in sub-Saharan Africa and Southeast Asia. There are five members of the *Plasmodium* genus known to infect humans, *Plasmodium falciparum* being most deadly. *P. falciparum* can not only evade the host immune system but can sequester and cyto-adhere to blood vessels leading to circulatory obstruction resulting in dysfunction of multiple organs including the brain. Understanding the mechanisms by which *P. falciparum* evades the immune response, and thus sustains further replication within the host, is key to identifying novel drug targets and abating the impact of malaria on global health.

To evade the host immune responses, the parasite switches the expression of specific exported proteins or antigens on the surface of infected red blood cells. The molecular mechanisms underlying ’antigenic variation’ have been extensively studied over the years with particular focus on the *var* gene family encoding *Plasmodium falciparum* erythrocyte membrane protein-1 (PfEMP-1). Expression of this gene family is thought to be regulated, at least partially, by epigenetic mechanisms [2, 3]. In eukaryotes, epigenetics involves various covalent modifications of nucleic acids and histone proteins which lead to changes in chromatin structure and gene expression. Similar mechanisms of gene regulation have been detected in *P. falciparum* with tight control of virulence genes organized in heterochromatin clusters with repressive histone mark H3K9me3 [4, 5, 6] localized at the nuclear periphery [7, 8].

PfEMP1 is exported to the plasma membrane of an infected red blood cell and mediates the adhesion of the infected erythrocytes to receptors on endothelial cells of the post capillary endothelium. Although there are approximately 60 members of the *var* gene family, only one *var* gene is expressed at any given time through a process known as mutually exclusive gene expression [9, 10]. To circumvent the host immune response, the parasite will switch the *var* gene expressed at a rate of 2-18% per generation preventing the recognition of parasitized cells by host antibodies [3, 11, 12]. During this switching process, it has been hypothesized that an intermediate state exists in which there is no dominantly expressed *var* gene and instead transcription is dispersed across several *var* genes before settling on a single dominant gene [13]. This was originally referred to as the “many” state and data supporting this model were recently obtained through transcriptomic analysis [14]. It is not known how parasites transition between these transcriptional states; however, this model implies a mechanism that unites the entire *var* gene family into a single network, potentially providing the basis for coordination of *var* gene expression within a large population of infected RBCs. Transcriptional dominance is complex and recent evidence in other models, including transcriptional regulation of odorant receptor genes, has identified possible non-coding mechanisms within mRNA that mediate the exclusive expression of a single gene within a large variable gene family [15].

Of the 60 *var* genes comprising the PfEMP1 family, *var2csa* is one of the most conserved between parasite strains and is the only variant whose transcription is upregulated in *P. falciparum*-parasitized erythrocytes in the placenta, causing clinical symptoms in the expectant mother and serious harm to the fetus [16, 17, 18]. Pregnancy associated malaria (PAM) results from sequestration of infected erythrocytes by binding chondroitin sulfate A (CSA) on the surface of syncytiotrophoblasts in the placental tissue [16]. Although expression on the surface of CSA-bound infected erythrocytes is exclusive to pregnant women, *var2csa* can be translationally repressed in infected erythrocytes while still being transcriptionally active at very low levels in non-pregnant populations, providing evidence that *var2csa* serves an additional, possible regulatory function [19, 20, 21, 22].

Although considerable progress has been made in elucidating the genetic and epigenetic basis for coordinated *var* gene switching, the hierarchy of importance in mechanisms controlling regulation remains unknown. Switching between different *var* genes is highly coordinated and imperative to parasite survival, but evidence suggests that some *var* genes may be more important than others and thus the rate of switching is variable depending on the location of the expressed *var* gene—subtelomeric versus central regions of the chromosomes [23, 13, 24, 25, 26]. Internal *var* genes are remarkably stable and transcriptional switching is uncommon compared to subtelomeric *var* genes which readily switch in the presence of environmental pressure [25, 26, 27]. Manipulation of H3K9me3 and H3K36me3 epigenetic marks results in upregulation of *var2csa* expression within a mixed population, regardless of previous *var* gene expression, suggesting that *var2csa* inhabits a prominent position within the *var* gene activation hierarchy [25, 28, 29]. Furthermore, when *var2csa* was deleted, *var* gene switching was disabled or greatly reduced, resulting in much more stable *var* gene expression over time [14]. This coincided with an overall reduction in the level of *var* transcripts, in particular the more dispersed transcriptional pattern observed in parasites transitioning through the hypothetical switching intermediate state. This implied that *var* gene expression had somehow become more tightly regulated, with fewer genes displaying even low-level transcriptional activity. However, how the loss of the *var2csa* locus led to this broader transcriptional phenotype is not known. Using a combination of transcriptomic, epigenetic and chromatin structure analyses we determined that *var2csa* deletion leads to a significant tightening of heterochromatin clusters throughout the genome, thus providing a likely explanation for reduced *var* transcription and disrupted transcriptional switching. These results confirm the importance of chromatin architecture during the process of *var* gene switching and provide a molecular basis for the phenotype resulting from deletion of the *var2csa* locus.

## RESULTS

### Deletion of *var2csa* alters *var* gene expression

To evaluate the effect of *var2csa* disruption on changes in overall *var* gene expression, we used Clustered Regularly Interspaced Short Palindromic Repeat/CRISPR-associated protein-9 (CRISPR/Cas9) targeted DNA double-strand breaks (DSBs) within the sub-telomeric region of chromosome 12 near the *var2csa* promoter (Fig 1A). The successfully edited cell line described in this manuscript as *var2csa*-deleted line (ΔV2) was previously shown to have undergone repair of the DSB through telomere healing by the addition of telomeric heptad repeat elements near the site of the break [30]. This resulted in the loss of the *var2csa* locus and surrounding region of the chromosome. Furthermore, although *var* gene transcription was not lost, deletion of *var2csa* was shown to greatly reduce *var* gene switching and resulted in a significant reduction in the *var* transcripts that make up a semi-conserved pattern of what were referred to as *var* “minor” transcripts [14]. These are low-level transcripts that come from a subset of *var* genes dispersed across the genome. How loss of *var2csa* led to a reduction in expression of transcripts coming from *var* genes located on other chromosomes was not clear, and any effects on members of other variant gene families was not determined.

**Figure 1.**
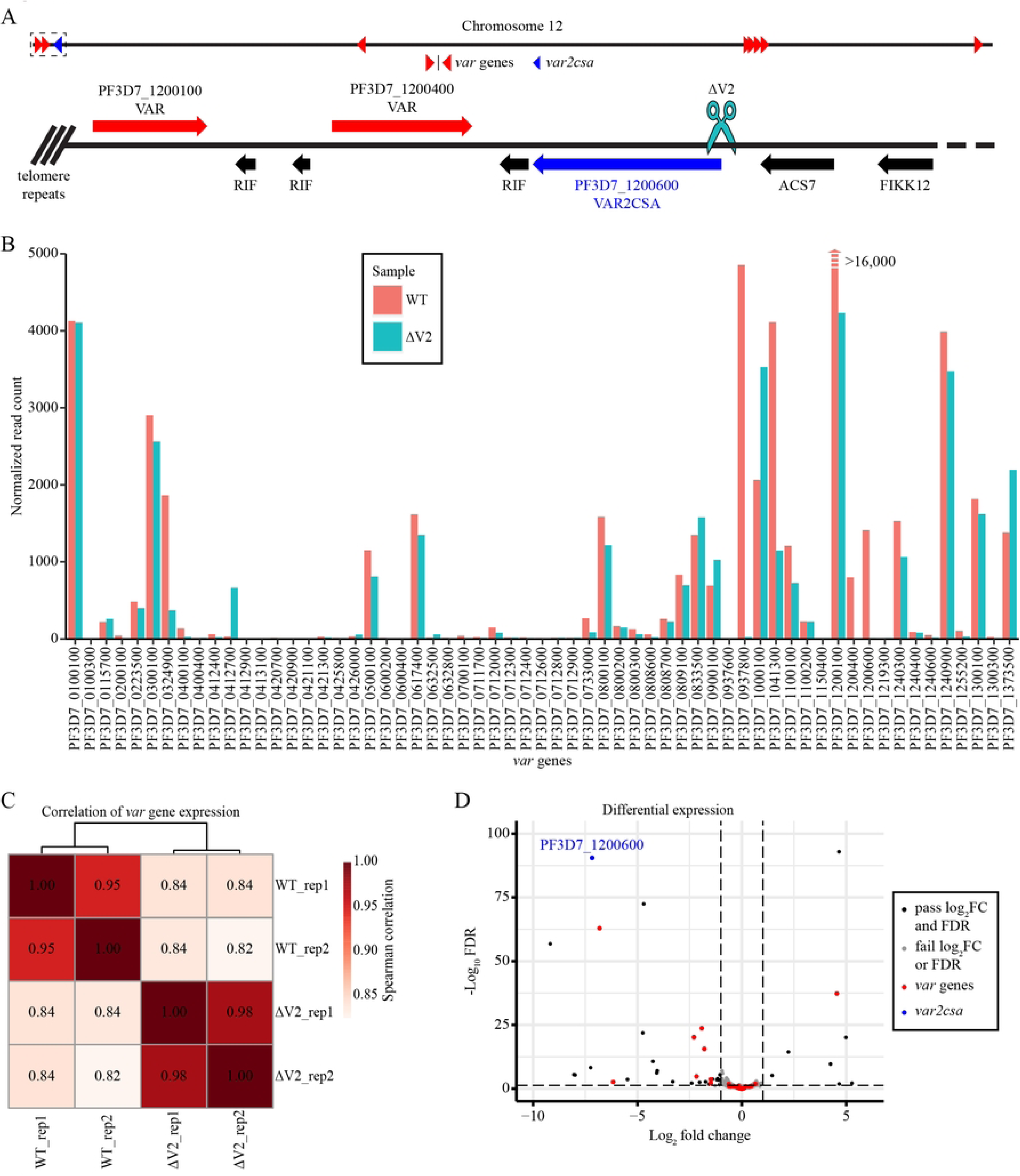
Overall *var* gene transcription decreased following a targeted DSB within the *var2csa* promoter region. (A) Location on chromosome 12 of the CRISPR/Cas9 targeted DSB within the *var2csa* promoter region. (B) Read count normalized *var* gene expression comparing the WT and ΔV2 cell lines. The figure was truncated to enhance visibility of other *var* genes due to significantly higher expression of PF3D7_1200100 in the WT cell line. (C) Spearman correlation for *var* gene expression in both replicates and cell lines. (D) Significant differentially expressed genes (log_2_FC > 1 and FDR < 0.05) with most *var* genes highlighted in red and *var2csa* in blue.

To further investigate the phenotype resulting from deletion of *var2csa*, we used RNA-seq to compare *var* gene expression of the ΔV2 line to an NF54 wild-type line (Fig 1B). Parasites and total RNA were collected at the trophozoite stage from two independent cultures [14]. Expression profile analysis revealed a strong correlation (*ρ_s_*> 0.98) between both replicates and samples (Fig S1A); however, the correlation for *var* gene expression was much weaker (*ρ_s_*≈ 0.82 to 0.84) between cell lines indicating that much of the transcriptional difference observed was due to changes in *var* gene expression (Fig 1C and S1A, B). Differential expression analysis revealed 51 significantly differentially expressed genes (FDR < 0.05 and log_2_FC > 1), with 10 upregulated and 41 downregulated (Fig 1D and S3 Table). Of the 51 significantly differentially expressed genes, 2 *var* genes display increased transcription (log_2_FC ≈ 4.61 and 0.77) and 10 *var* genes show decreased transcription (log_2_FC ≈ −0.74 to −7.66), including *var2csa* (log_2_FC ≈ −7.21). Furthermore, 11 of 14 *rifin* genes–an antigenic gene family found in close proximity to *var* gene clusters–are also downregulated. These significant differences are supported by the results of our gene ontology analysis, which shows significant enrichment of genes involved in antigenic variation for the downregulated genes (Fig S1C). These results confirm the total loss of *var2csa* expression following the introduction of the DSB, as well as the subsequent downregulation of several other *var* genes. Interestingly, the only highly upregulated *var* gene is located within an internal *var* gene cluster on chromosome 4, whereas all of the downregulated *var* genes are located in subtelomeric *var* gene clusters. This suggests that the knockout of *var2csa* results in epigenetic and chromatin structural changes that cause the downregulation of these subtelomerically located genes.

### Tighter *var* gene expression control by repressive histone marks

To better understand the role of epigenetics in *var* gene expression in our edited ΔV2 line, we assessed the distribution of repressive histone mark H3K9me3 and examined whether ΔV2 displays differentially bound heterochromatin due to *var2csa* deletion. We performed ChIP-seq experiments in duplicate in ΔV2 and NF54 WT lines using anti-H3K9me3 antibodies to identify whether the downregulated *var* genes strongly correlate with repressive histone marks. Our analysis confirmed an abundance of H3K9me3 on both subtelomeric and internal *var* gene regions in each chromosome. This result correlates well with previous evidence suggesting *var* gene-containing regions are tightly controlled in heterochromatin cluster(s) (Fig 2A) [7, 8]. H3K9me3 is characteristically absent from other regions of the genome, confirming that a sizable percentage of the genome is available for openly permissive gene expression with the exception of members of the variably expressed gene families, including *var* gene loci [31].

**Figure 2.**
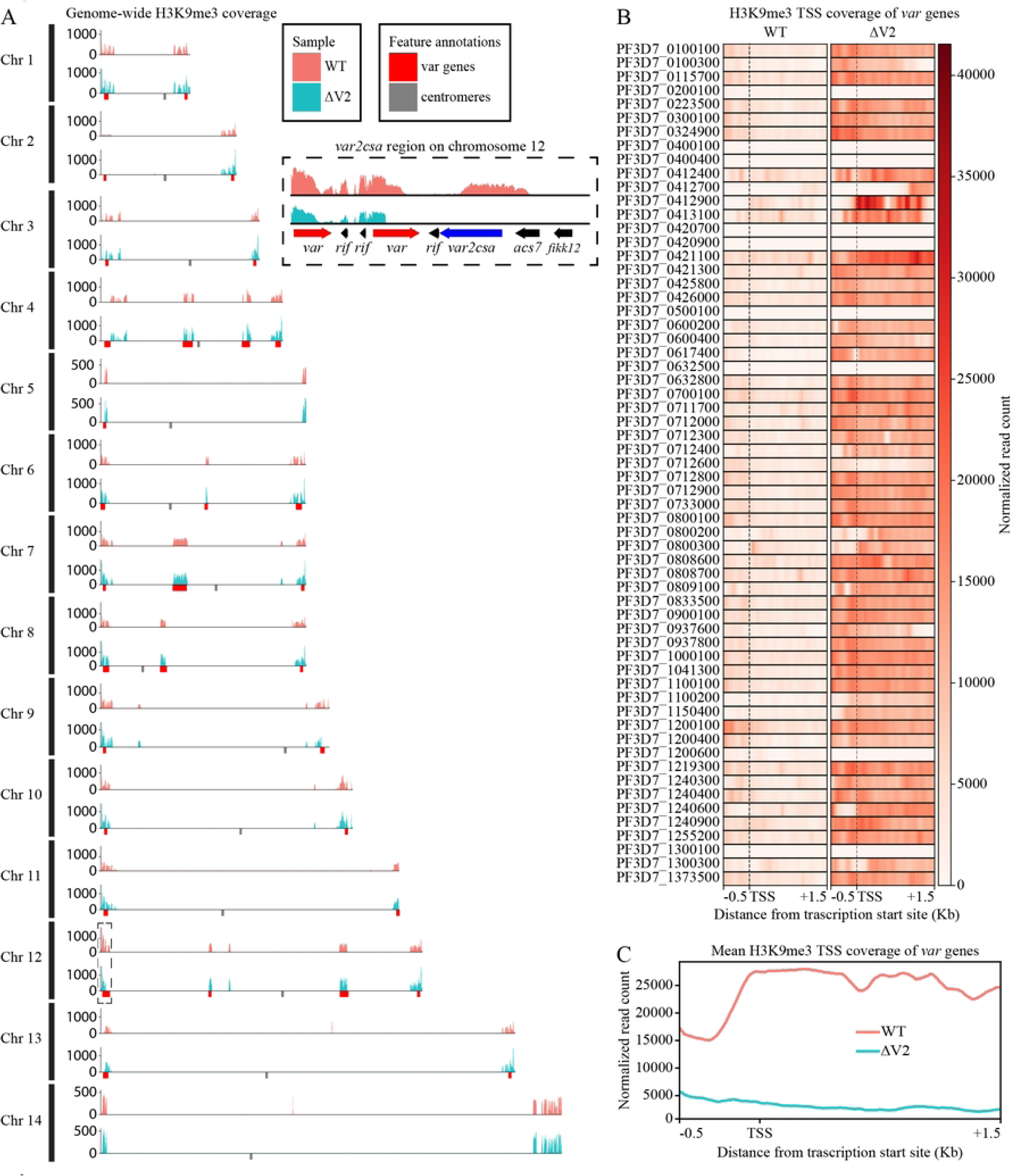
H3K9me3 is concentrated within *var* gene clusters and occupancy increased following *var2csa* disruption. (A) Genome-wide H3K9me3 coverage generated from per-million read count normalized coverage and input subtracted data. The small insert is the region surrounding *var2csa* within the subtelomeric region of chromosome 12. *Var* gene clusters are highlighted by red boxes along the x-axis and centromeres are denoted in gray. (B) Transcription start site (TSS) coverage of H3K9me3 for all *var* genes from 500 bp 5’ of each TSS and 1.5 kb 3’ of the TSS. (C) Profile plot of mean *var* TSS coverage showing the same region from (B) indicating much higher H3K9me3 coverage in ΔV2.

Peak calling analysis found 198 and 227 significant consensus peaks for the WT and ΔV2, respectively, covering less than 10% of the entire genome. Of the significantly called peaks located within 1 kb of a gene coding region, ∼25% of peaks are near *var* genes and an additional ∼36% are located near *rifin* genes (S4 Table and S5 Table). The transcription start site (TSS) coverage was ∼3-4x higher in *var* gene promoter regions and >5x higher across the gene body in ΔV2 compared to the wild type (Fig 2B, C). These data support our transcriptomic analysis and indicate that the down regulation of *var* genes following *var2csa* knockout is associated with an increase in heterochromatin-associated histone marks. Differential peak calling analysis found significant overlap between WT and ΔV2 peaks and only 12 differential binding sites (Fig S2). Of the 12 differentially called peaks, 4 reside within the telomeric or subtelomeric region of various chromosomes outside any gene coding regions, and 4 others are found in a subtelomeric region of chromosome 2 that is frequently spontaneously lost in cultured parasites (S6 Table) [32, 33]. This leaves only 4 broad coding regions that are differentially bound by H3K9me3 in ΔV2, suggesting that loss of *var* gene expression following *var2csa* knockout is not only due to changes in repressive histone marks.

### Structural changes to chromatin architecture linked to *var* expression

To confirm the link between epigenetics and chromatin structure, we then performed Hi-C experiments on tightly synchronized trophozoite stage parasites [34]. Hi-C libraries of two biological replicates for each sample (WT and ΔV2) were sequenced to a mean depth ∼56 million reads per replicate. After processing the libraries (aligning, pairing, mapping and quality filtering), we obtained ∼15-25 million valid interaction pairs per sample and binned the data at 10 kb resolution. Biological replicates display a strong stratum adjusted correlation (SCC > 0.94) with stronger similarity among ΔV2 samples compared to the WT control (Fig S3A). However, correlation between WT and ΔV2 samples was also strong (SCC > 0.80) suggesting concentrated structural differences in chromatin architecture. Due to strong correlation, biological replicates for each condition/genotype were merged to increase the data resolution for downstream analysis. Random sampling was then performed on the WT due to differences in sequencing depth and number of total valid interaction pairs. Interactions between telomeres and *var* gene-containing regions across all chromosomes, including internal *var* genes, are significantly higher than the background for both intrachromosomal and interchromosomal interactions in both WT and ΔV2 (Fig 3A and Fig S4, S5, S7A).

**Figure 3.**
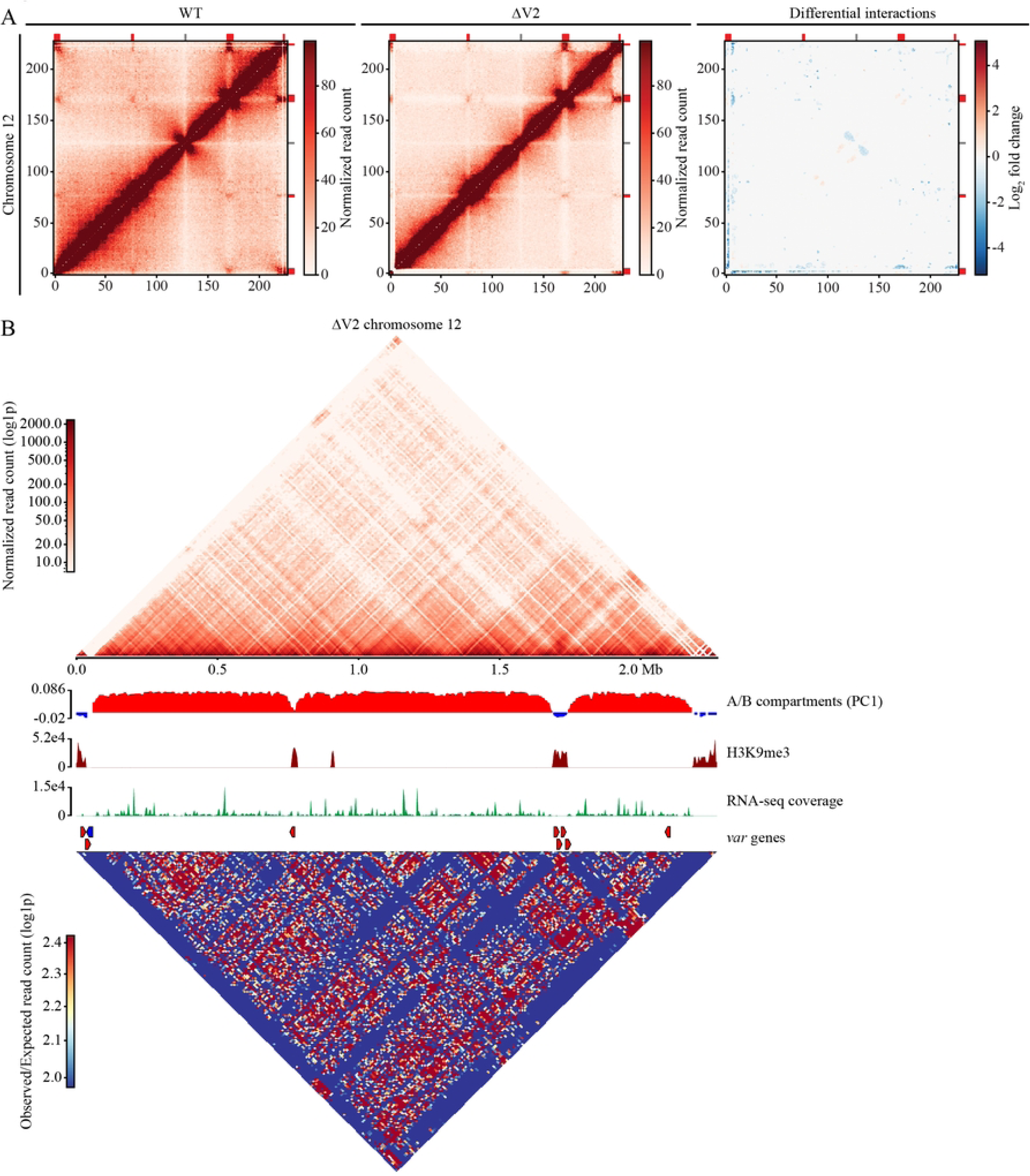
The targeted DSB resulted in a loss of chromatin interactions throughout most of the genome and compaction near *var* genes. (A) Hi-C contact count and differential interaction heatmaps for chromosome 12 binned at 10 kb resolution. The WT and ΔV2 interaction heatmaps are ICED normalized and the WT has been subsampled to the same number of reads as ΔV2. The color is scaled to the highest value in both datasets. (B) A series of plots for chromosome 12 of ΔV2, including log1p transformed ICED normalized read coverage, A/B compartment analysis, H3K9me3 coverage, RNA-seq transcript coverage, location of the *var* genes and plot of observed/expected interaction counts.

The difference in interaction counts between the background and subtelomeric regions in the wild type and ΔV2 is made even more apparent by the differential interaction analysis (q < 0.05), which revealed an increase in interchromosomal interactions between bins containing *var* genes and a general trend of decreased interactions between subtelomeric and internal *var* gene regions and other bins throughout the genome (Fig 3A and Fig S6). This indicates a chromatin restructuring where telomeres are moving away from the rest of the genome and condensing into even tighter heterochromatin structures. Decreases in chromatin interactions are also detected near the centromeres, resulting in a loss of centromeric A compartments in ΔV2 (Fig. 3B). The chromatin structure of the centromere is so distinct that principal component 1 (PC1) of the compartment analysis highlighted the B compartment surrounding the centromere in the WT rather than the dense heterochromatin *var* gene regions (Fig. S7B). The interactions near *P. falciparum* centromeres resemble the jet-like projections found in mammalian chromatin [35]. These chromatin jets are small regions of open chromatin that permeate nearby dense B compartments and extend perpendicularly, similar to the X-shaped structures near annotated centromeres and internal *var* gene clusters in the WT heatmaps (Fig. 3A, and Fig. S5). Hi-C analysis of chromatin jets in cohesin-depleted human lymphocytes shows substantial weakening of cohesin-dependent loop-extrusion and loss of chromatin jets [35]. Thus far no studies have been conducted on jet-like structures in *Plasmodium spp.*; therefore, additional experiments will be required to identify the breadth of molecular factors responsible for centromere and chromatin maintenance which result in these features. However, we provide compelling evidence to suggest that expression of *var2csa* is a key component necessary for the maintenance of the overall chromatin structure. Increased interaction in telomeric regions may lead to a loss of the jet-like structures observed near centromeres and internal *var* genes in *P. falciparum*.

To gain a more comprehensive view of the chromatin architecture and identify structural changes resulting from deletion of *var2csa*, we generated 3D models using Pastis [36]. The goal was to identify regions that show clear structural changes that would indicate chromatin remodeling resulting from deletion of *var2csa*, and perhaps the resulting recombination events, which would likely result in more compacted chromatin structure within *var* gene regions. Our models are consistent with previous work that utilized 3D chromatin modeling in that our models show centromeres and telomeres clustered into two separate distinct regions (Fig. 4) [37, 7, 38]. We see that compared to wildtype parasites, ΔV2 shows a more compacted chromatin structure involving *var* regions, with telomeres concentrated into closed structures with a decreased distance between telomeric *var* gene bins. Compaction of the telomeres in closed structures prevents access to transcriptional machinery and thus suppresses their expression, which is supported by the RNA-seq data showing downregulation of several subtelomeric *var* genes.

**Figure 4.**
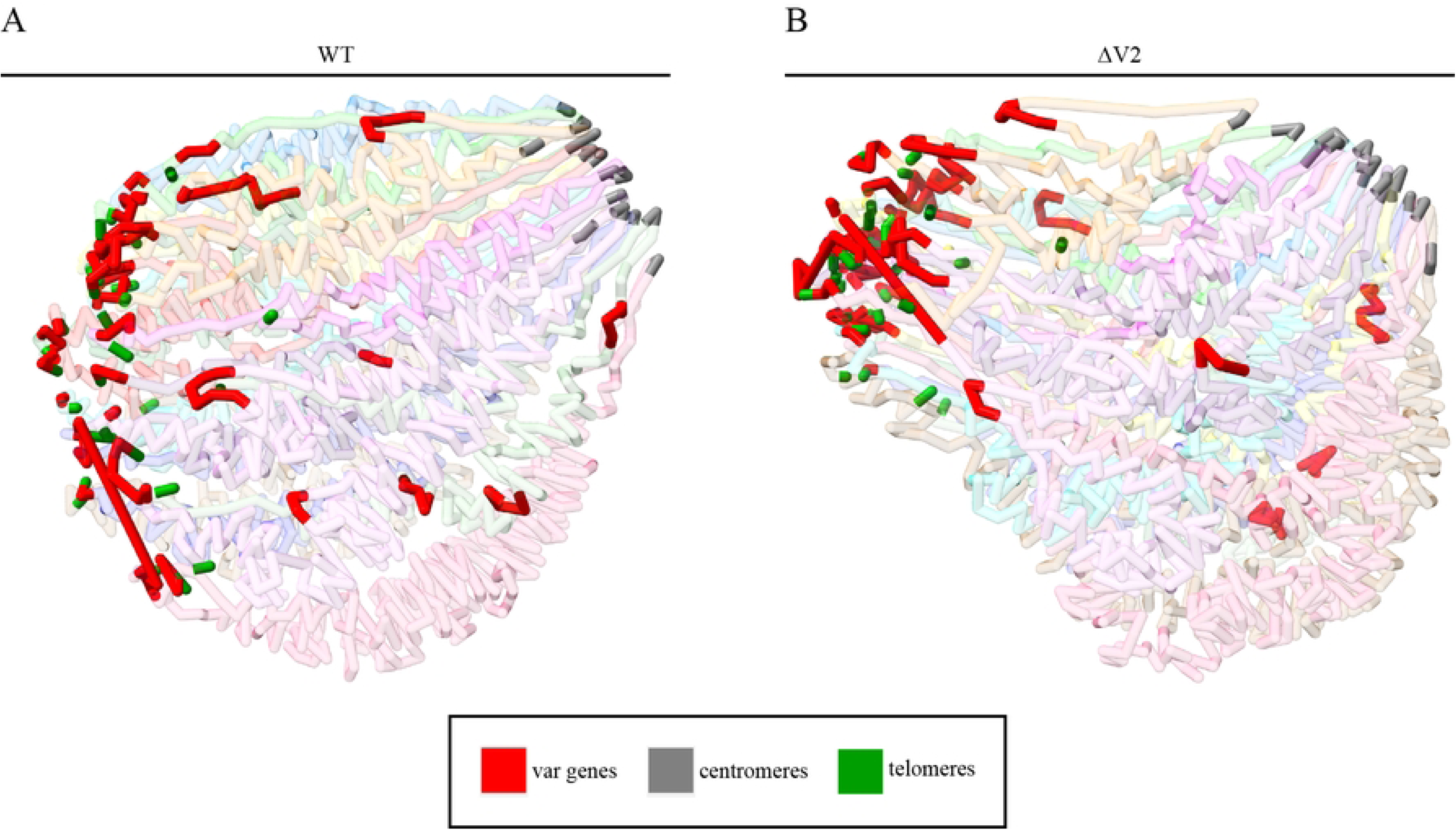
Genome-wide 3D models. (A) 3D chromatin model of the WT generated from ICED normalized contact count matrices. *Var* genes are highlighted in red, centromeres in gray and telomeres in green. (B) 3D model of ΔV2.

## DISCUSSION

Conservation of the highly variable *var* gene family within subtelomeric regions throughout the *P. falciparum* genome is imperative for adhesion of infected erythrocytes to receptors on endothelial cells. Expression of PfEMP1 and excessive cytoadherence of iRBCs within the microvasculature of various organs increases the severity of symptoms in malaria infections. Erythrocytes infected by VAR2CSA-expressing parasites bind to syncytiotrophoblasts within the placenta of pregnant women and can induce symptoms in previously immunocompetent women and cause significant damage to the fetus. The *var2csa* gene is unique among the *var* gene repertoire due to its specialized receptor preference and highly conserved sequence between parasite genomes, as well as being both transcriptionally and translationally regulated. Low-level transcription of *var2csa* in non-pregnant populations allows the parasite to be primed for proliferation within the placental tissue when infecting a pregnant woman while also not wasting energy on translation.

Maintenance of functional *var* genes aids the parasite in surviving the host immune response and avoiding splenic clearance through intermittent *var* gene switching, preventing immune recognition of the PfEMP1 surface proteins on iRBCs. Due to the importance of *var2csa* and maintenance of functional expression of PfEMP1, it is important to fully understand the underlying mechanisms controlling *var* gene expression and antigenic variation. The CRISPR-Cas9 targeted DSB within the promoter region of *var2csa* resulted in a complete loss of the locus. DNA repair via a cascade of non-homologous recombination events with the telomeres of other chromosomes produces chimeric *var* genes and ultimately supports parasite survival [30]. Previous work indicated that loss of *var2csa* reduced *var* transcriptional switching and led to a reduction in detectable *var* transcripts [14]. Our analysis confirmed a moderate downregulation of several *var* genes and other nearby subtelomeric genes. Differential expression analysis indicated that *var2csa* deletion affected transcription of subtelomeric genes almost exclusively, while transcription within the rest of the genome remained steady.

A model of promoter competition is frequently described as a possible mechanism underlying mutually exclusive expression within multicopy gene families in model organisms [39, 40]. We have similarly proposed that competition between *var* promoters could play an important role in selecting which *var* gene is active as well as contributing to *var* gene switching [41, 14]. The *var2csa* promoter appears to be unique in its competitiveness compared to other *var* promoters [21, 42], and thus its presence in the genome could serve to maintain expression levels of other *var* promoters within a range that is conducive to periodic switching. Its loss therefore could upset this balance, leading to reduced switching and lower levels of the *var* “minor” transcripts associated with *var* promoter activity. Consistent with this model, our Hi-C experiments showed that compared to the wildtype line, the *var2csa* deleted line displayed greater compaction of heterochromatin with increased interchromosomal interactions across subtelomeric *var* gene regions of all chromosomes. Compartment analysis also highlighted chromatin modifications near the centromere and loss of jet-like structures. The 3D chromatin models further emphasized the clear compaction of subtelomeric regions, which is in accordance with increased H3K9me3 silencing and downregulation of subtelomeric *var* gene expression. All together our data confirm the importance of chromatin structure and *var2csa* expression in *var* gene regulation. Detection of additional molecular factors, protein(s) or RNA(s), interacting with this locus will most likely be needed to fully comprehend at the mechanistic level sequential coding and non-coding components required for *var* gene expression.

## MATERIALS AND METHODS

### Samples and culturing

Parasite lines were NF54 isolates, one WT control line contributed by Megan G. Dowler and obtained through BEI Resources, NIAID, NIH: *Plasmodium falciparum* strain NF54 (Patient Line E), Catalog No. MRA-1000; and one line containing CRISPR/Cas9 targeted DSBs in a subtelomeric region of chromosome 12 (ΔV2). The parasites were maintained at ∼5-10% parasitemia in human erythrocytes at 5% hematocrit. Cultures were synchronized twice at the ring stage ∼8 hr apart using 5% sorbitol and then samples collected during the trophozoite stage at ∼6% parasitemia.

### RNA library preparation

Infected erythrocytes were treated with 0.15% saponin then flash frozen. Total RNA was extracted using TRIzol LS Reagent (Invitrogen) and chloroform and incubated overnight at −20°C in isopropanol. Samples were then treated with 4 units of DNase I (NEB) for 1 hr at 37°C. mRNA was then purified using NEBNext Poly(A) mRNA Magnetic Isolation Module (NEB, E7490) according to manufacturer’s instructions. Libraries were prepared using NEBNext Ultra Directional RNA Library Prep Kit (NEB, E7420) and PCR amplified (15 min at 37°C, 12 cycles of 30 sec at 98°C, 10 sec at 55°C, and 12 sec at 62°C, then 5 min at 62°C) with KAPA HiFi HotStart Ready Mix (Roche). Libraries were quantified by Bioanalyzer (Agilent) and sequenced using the NovaSeq 6000 (Illumina) and S4 300 flow cell for paired-end reads.

### RNA-seq data processing and differential expression analysis

Sample quality of the paired-end RNA-seq libraries was first assessed via FastQC (v.0.11.9) and the per-base sequence quality and content was used to determine read trimming length. The index adapters and an additional ∼12 bp were trimmed using Trimmomatic (v0.39) [43]. Paired and trimmed reads were then aligned to the *P. falciparum* 3D7 genome (PlasmoDB v58) using HISAT2 (v2.1.0) [44]. Non-properly mapped or paired reads and reads with a mapping quality score less than 30 were filtered using Samtools [45]. High quality properly paired and aligned reads were then mapped to protein coding genes using HTseq (v1.99.2) [46]. DESeq2 (v1.32.0) [47] was utilized to distinguish differentially expressed genes between the NF54 control line (WT) and the CRISPR-Cas9 modified line (ΔV2).

### ChIP-seq library preparation

Infected erythrocytes were collected in 1X PBS and lysed using 0.15% saponin. Chromatin was crosslinked using 1% formaldehyde then extracted with nuclear extraction buffer (10 mM HEPES, 10 mM KCl, 0.1 mM EDTA, 0.1 mM EGTA, 1 mM DTT, 2 mM AEBSF, 1X protease inhibitor and 1X phosphatase inhibitor) and homogenized via 26.5-gauge needle. DNA was sheared to an expected size of ∼200-700 bp using a Covaris S220 sonicator for 8 min with the following settings: 5% duty cycle, 140 PIP, 200 cycles per burst. Immunoprecipitation was performed with IGG and anti-H3K9me3 antibodies after aliquoting each sample for non-immunoprecipitated inputs. Immunoprecipitated DNA was extracted using phenol/chloroform, isoamyl (25:24:1). Libraries prepared using the KAPA Library Preparation Kit (KAPA Biosystems) and amplified using NEB indexed adapters with 12 PCR cycles (15 sec at 98°C, 30 sec at 55°C, and 30 sec at 62°C). Libraries were quantified by NanoDrop and Bioanalyzer (Agilent), then sequenced using the NovaSeq 6000 (Illumina) and S4 300 flow cell for paired-end reads.

### ChIP-seq data processing and peak calling

Paired-end ChIP-seq libraries were processed by FastQC (v0.11.9) for quality assessment. Indexes and low-quality reads were trimmed from the ends before pairing forward and reverse reads using Trimmomatic (v0.39) [43]. Bowtie2 [48] was then used to align paired reads to the *P. falciparum* 3D7 genome (PlasmoDB v58) with the “–very-sensitive” option to ensure the highest quality alignment. PCR duplicates were tagged using Picardtools (v2.26.11) [49]. Samtools [45] was utilized to filter, sort and index the aligned, de-duplicated reads with a mapping quality cutoff of 40 and keeping only mapped and properly paired reads. To remove background noise and analyze genome-wide coverage, reads were mapped to the genome at 1-bp resolution using Samtools and then counts-per-million normalized input reads were subtracted from H3K9me3 samples. Broad peak calling was performed using MACS3 (v3.0.0a7) [50] for peaks with FDR < 0.05. Coverage of the transcription start site (TSS) was evaluated with deeptools2 (v3.5.1) [51] by first using bamCompare to subtract input reads and then computeMatrix to map reads from 500 bp 5’ of the TSS to 1.5 kb 3’ of the TSS at 1-bp resolution. Differential peak calling was then performed using DiffBind (v3.2.7) [52].

### Hi-C library preparation

Parasite chromatin was crosslinked with 1.25% formaldehyde in warm 1X PBS. Cells were incubated in lysis buffer (10 mM Tris-HCl, pH 8.0, 10 mM NaCl, 2 mM AEBSF, 0.10% Igepal CA-360 (v/v), and 1X protease inhibitor cocktail) then homogenized with a 26-gauge needle. Crosslinked DNA was digested using MboI (NEB) restriction enzyme then 5’ overhangs were filled in by dNTPs with Biotin-14-dCTP (Invitrogen) using DNA Polymerase I and Large (Klenow) Fragment (NEB). Blunt-ends were ligated with 4000 units T4 DNA Ligase and chromatin de-crosslinked using de-crosslinking buffer (50 mM Tris-HCl at pH 8.0, 1% SDS, 1 mM EDTA, and 500 mM NaCl). Biotinylated DNA was sheared to 300-500 bp using a Covaris S220 (settings: 10% duty factor, 140 peak incident power, and 200 cycles per burst for 65 seconds) and biotinylated DNA fragments were pulled down using MyOne Streptavidin T1 beads (Invitrogen). End repair, A-tailing, and adapter ligation were all performed in lo-bind tubes using the KAPA library preparation kit (KAPA biosystems). Library was amplified using the HiFi HotStart ReadyMix (KAPA Biosystems) as well as the universal forward primer and barcoded reverse primer and incubated following the PCR program: 45 sec at 98°C, 12 cycles of 15 sec at 98°C, 30 sec at 55°C, 30 sec at 62°C and a final extension of 5 min at 62°C. The library was then purified using double-SPRI size selection, with 0.5Å∼ right-side selection (25 μl AMPure XP beads) and 1.0Å∼ left-side selection (25 μl AMPure XP beads). Libraries were quantified by NanoDrop (Thermo Scientific) and Bioanalyzer (Agilent), prior to multiplexing and sequencing in a 150-bp paired-end run on a NovaSeq 6000 (Illumina).

### Hi-C Data Analysis

Paired-end HiC library reads were processed (i.e., mapping, filtering, and pairing, and normalization) using the HiC-Pro suite [53] and the *P. falciparum* 3D7 genome (PlasmoDB v58) with mapping quality cutoff set to 30 and mapping at 10 kb resolution. Intra-chromosomal and inter-chromosomal ICED-normalized interaction matrices were interaction counts-per-million normalized before generating interaction heatmaps, with all contacts less than 2 bins apart set to 0 to enhance visualization. Differential interaction analysis was conducted at 10 kb resolution using Selfish while removing sparse regions [54]. HiCExplorer was used to identify and plot A/B compartments and O/E plots [55]. 3-dimensional chromatin remodeling was performed by generating 3-D coordinate matrices using Pastis [35] and then visualizing using ChimeraX [56].

## AUTHOR CONTRIBUTIONS

Conceptualization: Kirk Deitsch, Karine Le Roch

Data Curation: Todd Lenz

Formal analysis: Todd Lenz, Abhijit Chakraborty, Abbas Roayaei Ardakany, Xu Zhang, Ferhat Ay

Funding Acquisition: Karine Le Roch, Kirk Deitsch, Ferhat Ay. NIH/NIAID Grant # R21 AI142506-01 and R01 AI136511.

Investigation: Todd Lenz

Methodology: Xu Zhang, Kirk Deitsch

Project Administration: Karine Le Roch

Resources: Jacques Prudhomme

Software: Todd Lenz, Abhijit Chakraborty, Abbas Roayaei Ardakany, Ferhat Ay

Supervision: Karine Le Roch, Kirk Deitsch, Ferhat Ay

Validation: Todd Lenz, Karine Le Roch

Visualization: Todd Lenz

Writing – Original Draft Preparation: Todd Lenz, Karine Le Roch, Kirk Deitsch

## DATA AVAILABILITY

All raw sequence files (fastq) used to generate datasets for analyses within this study are available in the NCBI sequence read archive (SRA) under the project accession number PRJNA1069851. Any downstream files (e.g., BAM files and Hi-C matrices) are available upon request.

## ACKNOWLEDGEMENTS

This work was supported by the National Institutes of Allergy and Infectious Diseases and the National Institutes of Health [R01 AI136511, R21 AI142506 to K.L.R.]; and The University of California, Riverside |NIFA-Hatch-225935 to K.L.R.].

---

**Figure S1.**
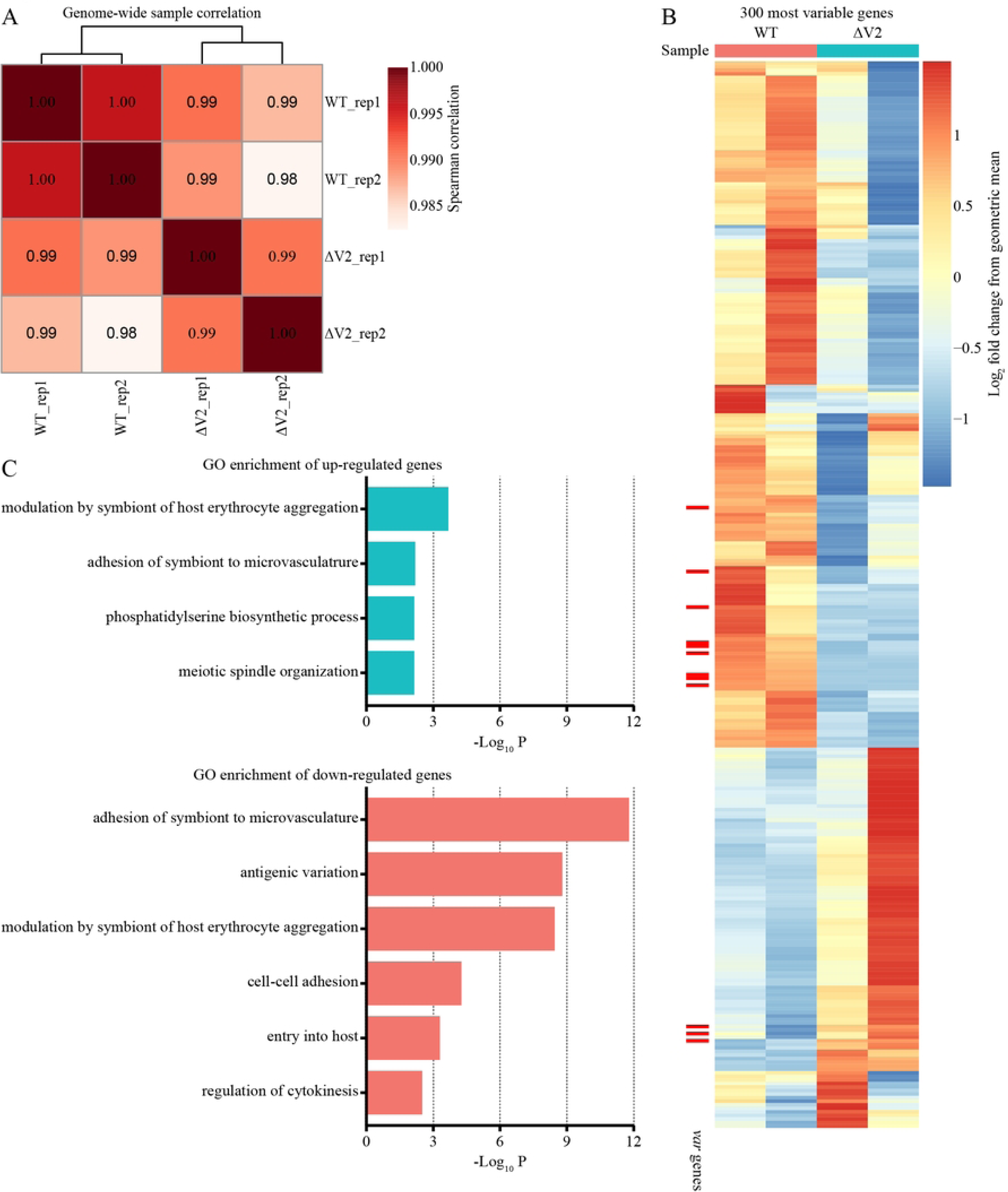
Differential expression analysis. (A) Spearman correlation for genome-wide expression in both replicates and cell lines. (B) Differential expression for the top 300 most variable genes among all samples with the *var* genes highlighted in red on the left side of the heatmap. (C) GO enrichment analysis for up-and downregulated genes in the differential expression dataset.

**Figure S2.**
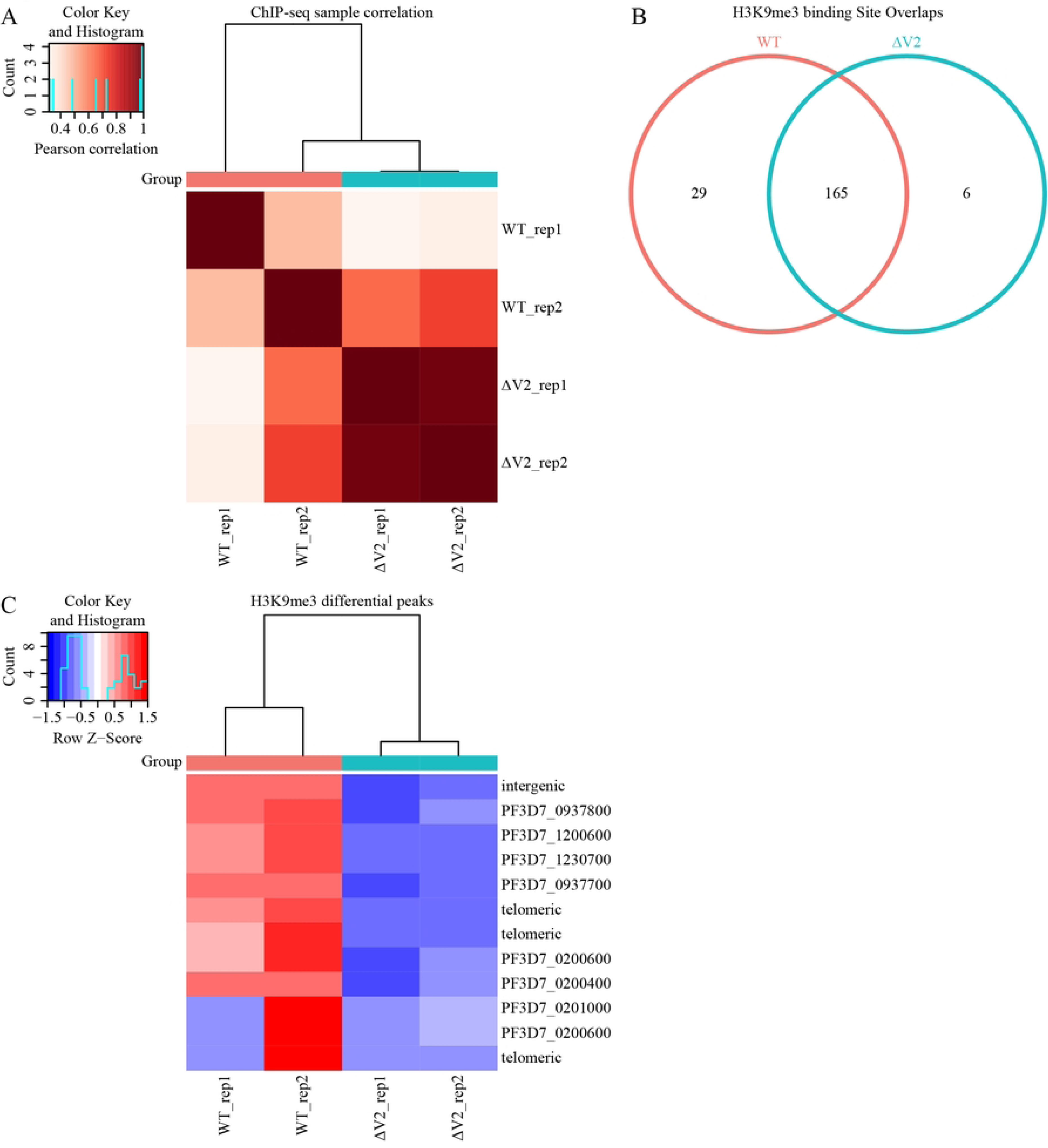
Differential peak calling between the WT and ΔV2 cell lines. (A) Pearson correlation of H3K9me3 peaks between all samples. (B) Number of unique and overlapping significant peaks for each cell line. (C) Deviation from mean read count in significant differential H3K9me3 binding sites. Differential peaks are mapped to genes within 1 kb of coding region.

**Figure S3.**
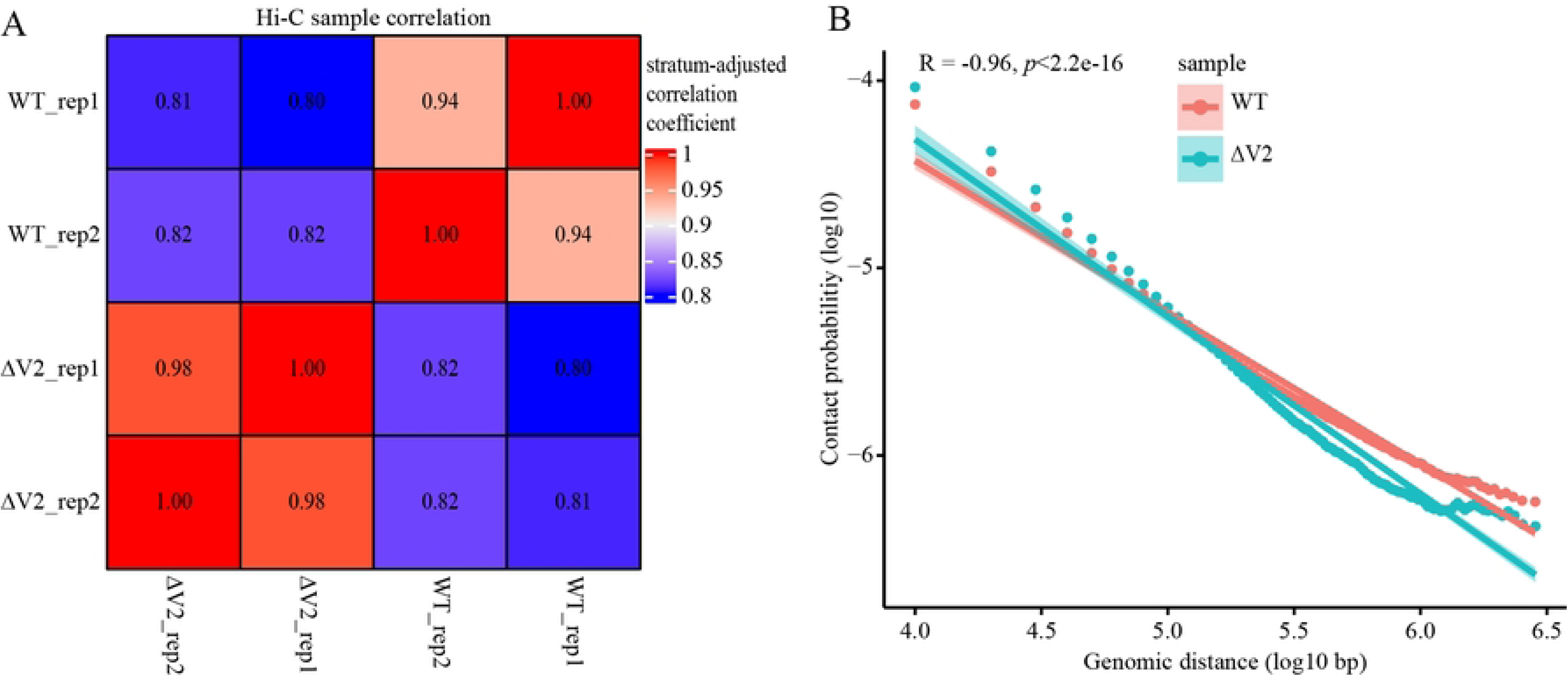
Hi-C correlation analysis. (A) Stratum-adjusted correlation between all samples used in the Hi-C analysis. (B) Negative log-linear relationship between genomic distance and contact probability.

**Figure S4-S5.**
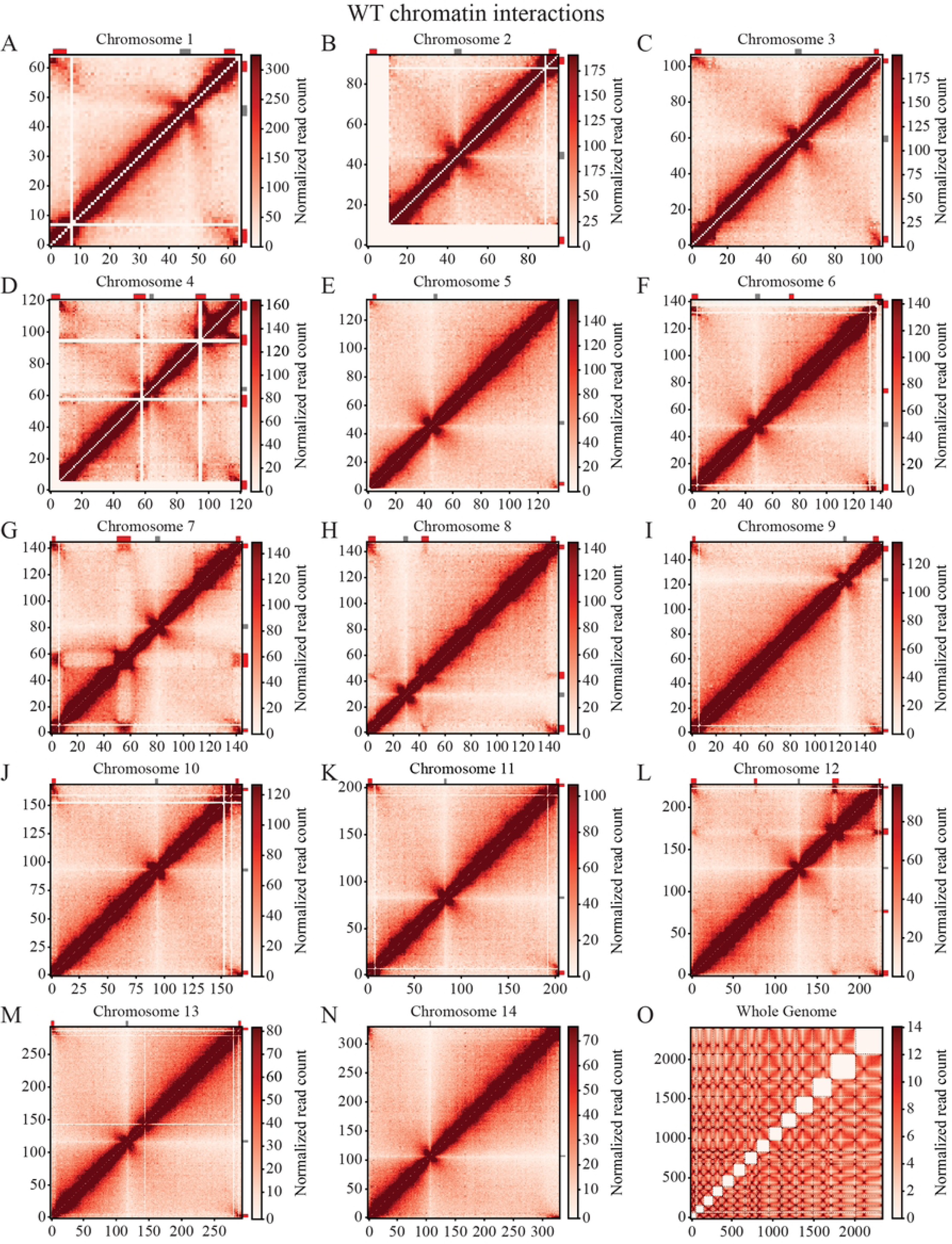
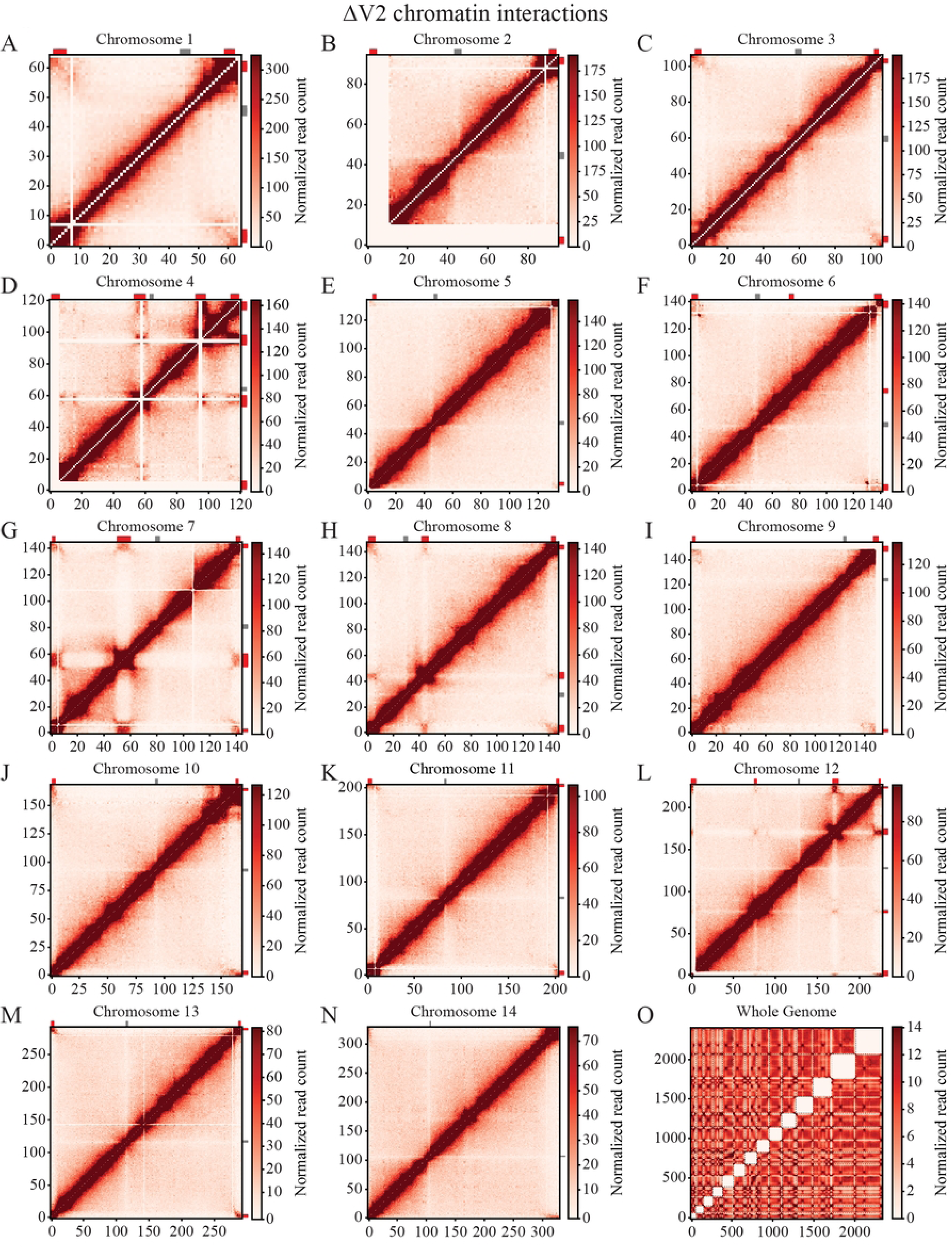
Hi-C interaction heatmaps for the WT (Fig. S4) and ΔV2 (Fig. S5) cell lines. Intrachromosomal interaction heatmaps (**A-N**) for each chromosome and an interchromosomal interaction heatmap (**O**) all binned at 10 kb resolution. All data is ICED normalized and WT sample is subsampled to the same read depth as ΔV2 with the color scaled to the highest value for each chromosome between both datasets. *Var* genes are indicated along the top and right side of each heatmap in red and the centromeres are shown in gray.

**Figure S6.**
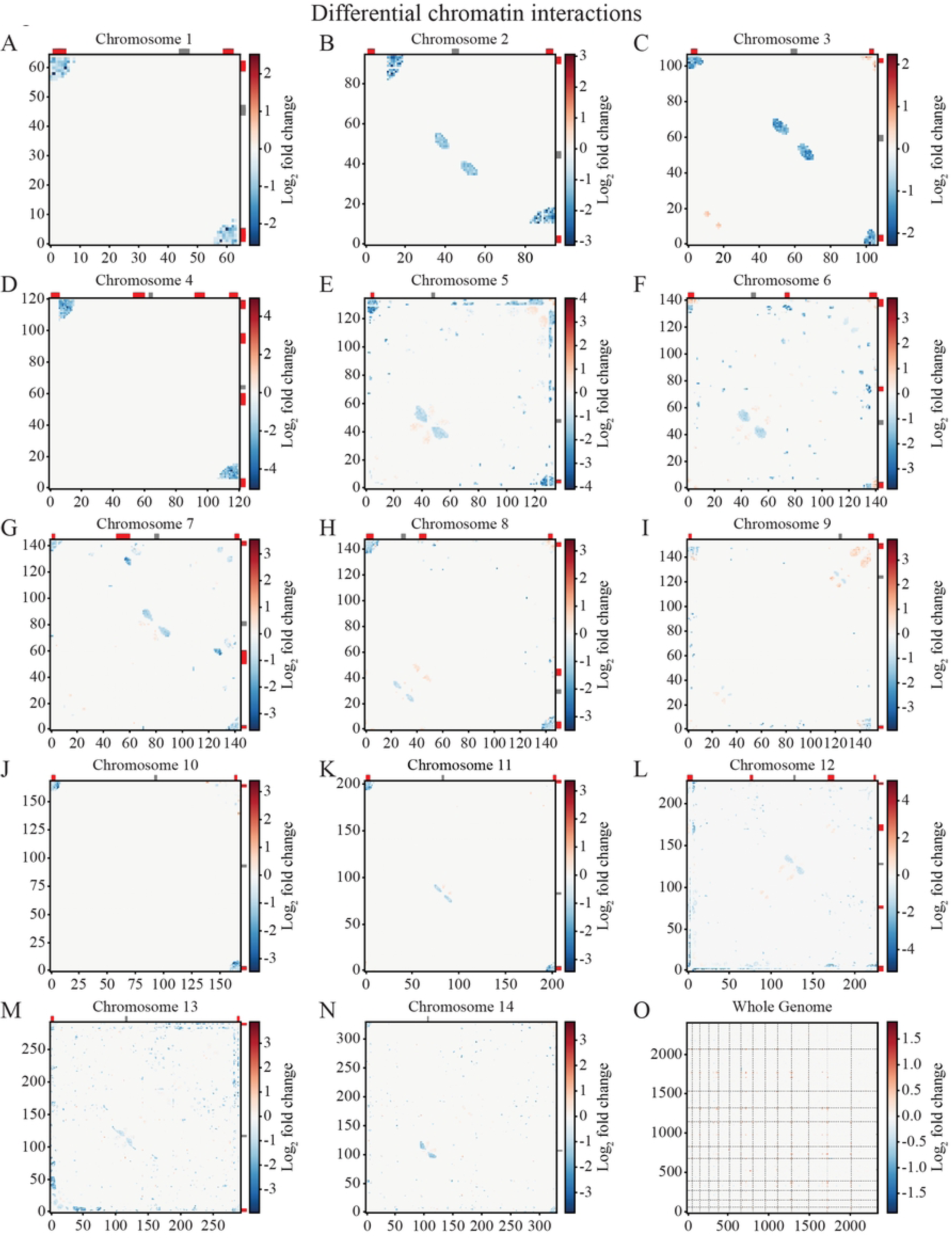
Differential Hi-C contact heatmaps. (A) Differential intra-and interchromosomal interaction heatmaps identifying regions with a positive (red) and negative (blue) log_2_ fold change in ΔV2 over WT. Bins containing *var* genes are indicated in red and centromeres in gray along the top and side of each heatmap.

**Figure S7.**
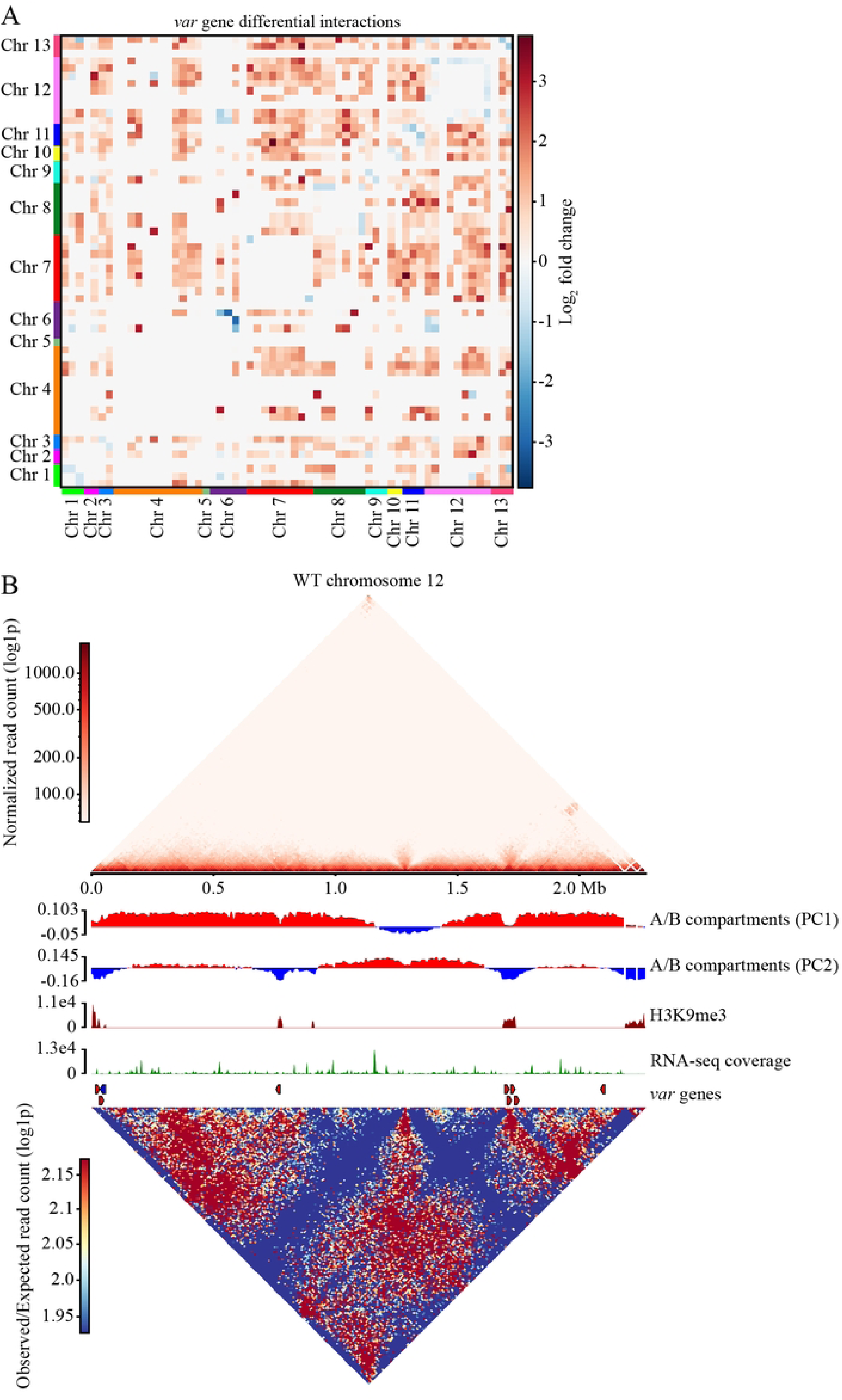
*Var* gene differential interactions and compartment analysis of the WT. (A) Differential interactions for 10 kb bins containing *var* genes. *Var* genes that fall within two bins are mapped to the bin containing the 5’ end and promoter region of the gene. (B) A series of plots for chromosome 12 of the WT, including log1p transformed ICED normalized read coverage, PC1 and PC2 of the A/B compartment analysis, H3K9me3 coverage, RNA-seq transcript coverage, location of the *var* genes and plot of observed/expected interaction counts.

**S1 Table. Sample data.** Table listing all samples for each experiment and the raw read count contained within the fastq files. The aligned/filtered read count is the number of valid reads contained within the output files following genome alignment and filtering.

**S2 Table. RNA-seq mapped reads.** Number of reads mapped to each gene within the *P. falciparum* genome for each sample in the RNA-seq experiment. Reads counts are not normalized.

**S3 Table. RNA-seq differential expression analysis.** Data from the differential expression analysis of ΔV2 versus WT. WT was set as the control; therefore, a positive log_2_ fold change indicates a gene that is more highly expressed in ΔV2.

**S4 Table. WT ChIP-seq peak calling.** MACS2 broad peak calling analysis of the WT samples using a q-value cutoff of 0.05. Peaks were subsequently mapped to the nearest gene if the peak is within 1 kb of the coding region of the gene.

**S5 Table. ΔV2 ChIP-seq peak calling.** MACS2 broad peak calling analysis of the ΔV2 samples using a q-value cutoff of 0.05. Peaks were subsequently mapped to the nearest gene if the peak is within 1 kb of the coding region.

**S6 Table. ChIP-seq differential peak calling analysis.** Results from the DiffBind differential peak calling of ΔV2 versus WT. A positive log_2_ fold change indicates a peak that contains elevated H3K9me3 in ΔV2. Peaks were mapped to the nearest gene if the differential peak is within 1 kb of the coding region.

## Notes

### Competing Interest Statement

The authors have declared no competing interest.

## REFERENCES

1. WHO. World malaria report 2023. Geneva: World Health Organization; 2023.

2. Baruch DI, Pasloske BL, Singh HB, et al. Cloning the *P. falciparum* gene encoding PfEMP1, a malarial variant antigen and adherence receptor on the surface of parasitized human erythrocytes. Cell. 1995;82(1):77–87. doi:10.1016/0092-8674(95)90054-3

3. Smith JD, Chitnis CE, Craig AG, et al. Switches in expression of *Plasmodium falciparum var* genes correlate with changes in antigenic and cytoadherent phenotypes of infected erythrocytes. Cell. 1995;82(1):101–110. doi:10.1016/0092-8674(95)90056-x

4. Trelle MB, Salcedo-Amaya AM, Cohen AM, Stunnenberg HG, Jensen ON. Global histone analysis by mass spectrometry reveals a high content of acetylated lysine residues in the malaria parasite *Plasmodium falciparum*. J Proteome Res. 2009;8(7):3439–3450. doi:10.1021/pr9000898

5. Jiang L, Mu J, Zhang Q, et al. PfSETvs methylation of histone H3K36 represses virulence genes in *Plasmodium falciparum*. Nature. 2013;499(7457):223-227. doi:10.1038/nature12361

6. Salcedo-Amaya AM, van Driel MA, Alako BT, et al. Dynamic histone H3 epigenome marking during the intraerythrocytic cycle of *Plasmodium falciparum*. Proc Natl Acad Sci U S A. 2009;106(24):9655–9660. doi:10.1073/pnas.0902515106

7. Bunnik EM, Cook KB, Varoquaux N, et al. Changes in genome organization of parasite-specific gene families during the *Plasmodium* transmission stages. Nat Commun. 2018;9(1):1910. Published 2018 May 15. doi:10.1038/s41467-018-04295-5

8. Bunnik EM, Venkat A, Shao J, et al. Comparative 3D genome organization in apicomplexan parasites. Proc Natl Acad Sci U S A. 2019;116(8):3183–3192. doi:10.1073/pnas.1810815116

9. Scherf A, Hernandez-Rivas R, Buffet P, et al. Antigenic variation in malaria: in situ switching, relaxed and mutually exclusive transcription of *var* genes during intra-erythrocytic development in *Plasmodium falciparum*. EMBO J. 1998;17(18):5418–5426. doi:10.1093/emboj/17.18.5418

10. Dzikowski R, Frank M, Deitsch K. Mutually exclusive expression of virulence genes by malaria parasites is regulated independently of antigen production. PLoS Pathog. 2006;2(3):e22. doi:10.1371/journal.ppat.0020022

11. Roberts DJ, Craig AG, Berendt AR, et al. Rapid switching to multiple antigenic and adhesive phenotypes in malaria. Nature. 1992;357(6380):689-692. doi:10.1038/357689a0

12. Gatton ML, Peters JM, Fowler EV, Cheng Q. Switching rates of *Plasmodium falciparum var* genes: faster than we thought?. Trends Parasitol. 2003;19(5):202–208. doi:10.1016/s1471-4922(03)00067-9

13. Recker M, Buckee CO, Serazin A, et al. Antigenic variation in *Plasmodium falciparum* malaria involves a highly structured switching pattern. PLoS Pathog. 2011;7(3):e1001306. doi:10.1371/journal.ppat.1001306

14. Zhang X, Florini F, Visone JE, et al. A coordinated transcriptional switching network mediates antigenic variation of human malaria parasites. Elife. 2022;11:e83840. Published 2022 Dec 14. doi:10.7554/eLife.83840

15. Pourmorady AD, Bashkirova EV, Chiariello AM, et al. RNA-mediated symmetry breaking enables singular olfactory receptor choice. Nature. 2024;625(7993):181-188. doi:10.1038/s41586-023-06845-4

16. Fried M, Duffy PE. Adherence of *Plasmodium falciparum* to chondroitin sulfate A in the human placenta. Science. 1996;272(5267):1502-1504. doi:10.1126/science.272.5267.1502

17. Salanti A, Dahlbäck M, Turner L, et al. Evidence for the involvement of VAR2CSA in pregnancy-associated malaria. J Exp Med. 2004;200(9):1197–1203. doi:10.1084/jem.20041579

18. Trimnell AR, Kraemer SM, Mukherjee S, et al. Global genetic diversity and evolution of *var* genes associated with placental and severe childhood malaria. Mol Biochem Parasitol. 2006;148(2):169–180. doi:10.1016/j.molbiopara.2006.03.012

19. Duffy MF, Caragounis A, Noviyanti R, et al. Transcribed *var* genes associated with placental malaria in Malawian women. Infect Immun. 2006;74(8):4875–4883. doi:10.1128/IAI.01978-05

20. Lavstsen T, Magistrado P, Hermsen CC, et al. Expression of *Plasmodium falciparum* erythrocyte membrane protein 1 in experimentally infected humans. Malar J. 2005;4:21. Published 2005 Apr 27. doi:10.1186/1475-2875-4-21

21. Mok BW, Ribacke U, Rasti N, et al. Default Pathway of *var2csa* switching and translational repression in *Plasmodium falciparum*. PLoS One. 2008;3(4):e1982. Published 2008 Apr 23. doi:10.1371/journal.pone.0001982

22. Amulic B, Salanti A, Lavstsen T, Nielsen MA, Deitsch KW. An upstream open reading frame controls translation of *var2csa*, a gene implicated in placental malaria. PLoS Pathog. 2009;5(1):e1000256. doi:10.1371/journal.ppat.1000256

23. Horrocks P, Pinches R, Christodoulou Z, Kyes SA, Newbold CI. Variable *var* transition rates underlie antigenic variation in malaria. Proc Natl Acad Sci U S A. 2004;101(30):11129–11134. doi:10.1073/pnas.0402347101

24. Noble R, Christodoulou Z, Kyes S, Pinches R, Newbold CI, Recker M. The antigenic switching network of *Plasmodium falciparum* and its implications for the immuno-epidemiology of malaria. Elife. 2013;2:e01074. Published 2013 Sep 17. doi:10.7554/eLife.01074

25. Ukaegbu UE, Zhang X, Heinberg AR, Wele M, Chen Q, Deitsch KW. A Unique Virulence Gene Occupies a Principal Position in Immune Evasion by the Malaria Parasite *Plasmodium falciparum*. PLoS Genet. 2015;11(5):e1005234. Published 2015 May 19. doi:10.1371/journal.pgen.1005234

26. Frank M, Dzikowski R, Amulic B, Deitsch K. Variable switching rates of malaria virulence genes are associated with chromosomal position. Mol Microbiol. 2007;64(6):1486–1498. doi:10.1111/j.1365-2958.2007.05736.x

27. Bachmann A, Predehl S, May J, et al. Highly co-ordinated *var* gene expression and switching in clinical *Plasmodium falciparum* isolates from non-immune malaria patients. Cell Microbiol. 2011;13(9):1397–1409. doi:10.1111/j.1462-5822.2011.01629.x

28. Sander AF, Salanti A, Lavstsen T, et al. Multiple *var2csa*-type PfEMP1 genes located at different chromosomal loci occur in many *Plasmodium falciparum* isolates. PLoS One. 2009;4(8):e6667. Published 2009 Aug 19. doi:10.1371/journal.pone.0006667

29. Brolin KJ, Ribacke U, Nilsson S, et al. Simultaneous transcription of duplicated *var2csa* gene copies in individual *Plasmodium falciparum* parasites. Genome Biol. 2009;10(10):R117. doi:10.1186/gb-2009-10-10-r117

30. Zhang X, Alexander N, Leonardi I, Mason C, Kirkman LA, Deitsch KW. Rapid antigen diversification through mitotic recombination in the human malaria parasite *Plasmodium falciparum*. PLoS Biol. 2019;17(5):e3000271. Published 2019 May 13. doi:10.1371/journal.pbio.3000271

31. Michel-Todó L, Bancells C, Casas-Vila N, et al. Patterns of Heterochromatin Transitions Linked to Changes in the Expression of *Plasmodium falciparum* Clonally Variant Genes. Microbiol Spectr. 2023;11(1):e0304922. doi:10.1128/spectrum.03049-22

32. Horrocks P, Kyes S, Pinches R, Christodoulou Z, Newbold C. Transcription of subtelomerically located *var* gene variant in *Plasmodium falciparum* appears to require the truncation of an adjacent *var* gene. Mol Biochem Parasitol. 2004;134(2):193–199. doi:10.1016/j.molbiopara.2003.11.016

33. Figueiredo LM, Freitas-Junior LH, Bottius E, Olivo-Marin JC, Scherf A. A central role for *Plasmodium falciparum* subtelomeric regions in spatial positioning and telomere length regulation. EMBO J. 2002;21(4):815–824. doi:10.1093/emboj/21.4.815

34. Rao SS, Huntley MH, Durand NC, et al. A 3D map of the human genome at kilobase resolution reveals principles of chromatin looping [published correction appears in *Cell*. 2015 Jul 30;162(3):687-8]. Cell. 2014;159(7):1665–1680. doi:10.1016/j.cell.2014.11.021

35. Guo Y, Al-Jibury E, Garcia-Millan R, et al. Chromatin jets define the properties of cohesin-driven in vivo loop extrusion. Mol Cell. 2022;82(20):3769–3780.e5. doi:10.1016/j.molcel.2022.09.003

36. Varoquaux N, Ay F, Noble WS, Vert JP. A statistical approach for inferring the 3D structure of the genome. Bioinformatics. 2014;30(12):i26–i33. doi:10.1093/bioinformatics/btu268

37. Bunnik EM, Polishko A, Prudhomme J, et al. DNA-encoded nucleosome occupancy is associated with transcription levels in the human malaria parasite *Plasmodium falciparum*. BMC Genomics. 2014;15(1):347. Published 2014 May 8. doi:10.1186/1471-2164-15-347

38. Ay F, Bunnik EM, Varoquaux N, et al. Three-dimensional modeling of the *P. falciparum* genome during the erythrocytic cycle reveals a strong connection between genome architecture and gene expression. Genome Res. 2014;24(6):974–988. doi:10.1101/gr.169417.113

39. Hutchinson S, Foulon S, Crouzols A, et al. The establishment of variant surface glycoprotein monoallelic expression revealed by single-cell RNA-seq of Trypanosoma brucei in the tsetse fly salivary glands. PLoS Pathog. 2021;17(9):e1009904. Published 2021 Sep 20. doi:10.1371/journal.ppat.1009904

40. Constância M, Pickard B, Kelsey G, Reik W. Imprinting mechanisms. Genome Res. 1998;8(9):881-900. doi:10.1101/gr.8.9.881

41. Dzikowski R, Deitsch KW. Active transcription is required for maintenance of epigenetic memory in the malaria parasite *Plasmodium falciparum*. J Mol Biol. 2008;382(2):288–297. doi:10.1016/j.jmb.2008.07.015

42. Schneider VM, Visone JE, Harris CT, et al. The human malaria parasite *Plasmodium falciparum* can sense environmental changes and respond by antigenic switching. Proc Natl Acad Sci U S A. 2023;120(17):e2302152120. doi:10.1073/pnas.2302152120

43. Bolger AM, Lohse M, Usadel B. Trimmomatic: a flexible trimmer for Illumina sequence data. Bioinformatics. 2014;30(15):2114–2120. doi:10.1093/bioinformatics/btu170

44. Kim D, Langmead B, Salzberg SL. HISAT: a fast spliced aligner with low memory requirements. Nat Methods. 2015;12(4):357–360. doi:10.1038/nmeth.3317

45. Li H, Handsaker B, Wysoker A, et al. The Sequence Alignment/Map format and SAMtools. Bioinformatics. 2009;25(16):2078–2079. doi:10.1093/bioinformatics/btp352

46. Anders S, Pyl PT, Huber W. HTSeq--a Python framework to work with high-throughput sequencing data. Bioinformatics. 2015;31(2):166–169. doi:10.1093/bioinformatics/btu638

47. Love MI, Huber W, Anders S. Moderated estimation of fold change and dispersion for RNA-seq data with DESeq2. Genome Biol. 2014;15(12):550. doi:10.1186/s13059-014-0550-8

48. Langmead B, Salzberg SL. Fast gapped-read alignment with Bowtie 2. Nat Methods. 2012;9(4):357–359. Published 2012 Mar 4. doi:10.1038/nmeth.1923

49. Broad Institute. Picard toolkit. Broad Institute, GitHub repository. 2019. https://broadinstitute.github.io/picard/

50. Zhang Y, Liu T, Meyer CA, et al. Model-based analysis of ChIP-Seq (MACS). Genome Biol. 2008;9(9):R137. doi:10.1186/gb-2008-9-9-r137

51. Ramírez F, Ryan DP, Grüning B, et al. deepTools2: a next generation web server for deep-sequencing data analysis. Nucleic Acids Res. 2016;44(W1):W160–W165. doi:10.1093/nar/gkw257

52. Stark R, Brown G. DiffBind: differential binding analysis of ChIP-Seq peak data. 2011. http://bioconductor.org/packages/release/bioc/vignettes/DiffBind/inst/doc/DiffBind.pdf

53. Servant N, Varoquaux N, Lajoie BR, et al. HiC-Pro: an optimized and flexible pipeline for Hi-C data processing. Genome Biol. 2015;16:259. Published 2015 Dec 1. doi:10.1186/s13059-015-0831-x

54. Ardakany AR, Ay F, Lonardi S. Selfish: discovery of differential chromatin interactions via a self-similarity measure. Bioinformatics. 2019;35(14):i145–i153. doi:10.1093/bioinformatics/btz362

55. Ramírez F, Bhardwaj V, Arrigoni L, et al. High-resolution TADs reveal DNA sequences underlying genome organization in flies. Nat Commun. 2018;9(1):189. Published 2018 Jan 15. doi:10.1038/s41467-017-02525-w

56. Goddard TD, Huang CC, Meng EC, et al. UCSF ChimeraX: Meeting modern challenges in visualization and analysis. Protein Sci. 2018;27(1):14–25. doi:10.1002/pro.3235

